# Normalized Dynamic Time Warping Increases Sensitivity in Differentiating Functional Network Connectivity in Schizophrenia

**DOI:** 10.1101/2024.10.31.621415

**Authors:** S-L. Wiafe, S. Kinsey, A. Iraji, Robyn Miller, V. D. Calhoun

## Abstract

Our study advances the application of dynamic time warping (DTW) as a functional connectivity measure by introducing a normalization technique which enhances the detection of schizophrenia effects in comparison to both standard DTW and traditional correlation methods. By rigorously examining the statistical validity of DTW and our proposed normalized DTW measure, we show that it effectively captures interdependencies between fMRI signals beyond linear correlation, offering a more robust, complementary and informative approach to functional connectivity analysis. Through comprehensive evaluations, we demonstrate that normalized DTW is more sensitive to differences in functional brain network connections between schizophrenia and controls, highlighting its potential to provide deeper insight into clinical research.

**Clinical Relevance:** This study enhances our understanding of the functional specificity of schizophrenia by emphasizing the importance of nonlinear relationships through the introduction of a normalization technique for the DTW metric.

## I. Introduction

Recent advancements in functional magnetic resonance imaging (fMRI) and functional network connectivity (FNC) analysis have highlighted the superiority of dynamic time warping (DTW) over traditional correlation methods in aligning brain network signals and gauging the similarity of their time courses [1, 2]. DTW has demonstrated greater robustness to global signal regression and noise, enhanced sensitivity to group differences such as gender and autism spectrum disorder, and improved test-retest reliability [1, 2]. Moreover, we highlight *warp elasticity* as a time-resolved DTW-based method that captures the stretching and shrinking dynamics of fMRI brain network signals [3, 4], demonstrating the potential of DTW to provide new insights into functional brain organization.

DTW computes the cumulative distance between two time-series by allowing non-linear alignments, but the optimal alignment, warping path, can vary in length across signals. Because some signals require more alignment steps, DTW distances may be inflated or deflated simply by virtue of having longer or shorter warping paths. In group-level comparisons, this variability can confound interpretation by mixing genuine time-course dissimilarities with the “penalty” of a longer path. To address this issue, we propose a normalized DTW (nDTW) metric that divides DTW distance by warping path length, providing more comparable results across different signals. Furthermore, we demonstrate the statistical significance of both DTW and nDTW in capturing interdependencies between fMRI signals beyond linear correlations. Moreover, we replicate DTW’s sensitivity to group differences, showing that both DTW and nDTW outperform correlation in identifying group differences, particularly in distinguishing between controls (CN) and individuals with schizophrenia (SZ).

## II. Methods

### A. Dynamic time warping

Dynamic time warping (DTW) is a technique for measuring similarity between two signals that may vary in time or speed by minimizing a cumulative distance metric between corresponding points in the time series [5, 6]. The DTW algorithm calculates a distance cost matrix (CM) that captures the cumulative alignment cost of sequence elements and identifies the optimal path, or warping function, to minimize the distance between them.

To ensure meaningful alignment, a window size constraint often limits the maximum allowable warping between indices in the series [6, 7]. For discrete signals *x* and *y* of lengths *N* and *M* respectively, the distance cost matrix, *CM*, that within the allowable window, *w*, can be defined as:

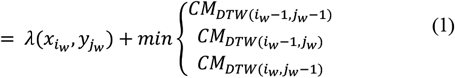

Where λ is a distance metric, 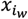 and 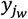 are timepoints within the allowable window *w* for *x* and *y* respectively. In DTW for fMRI functional connectivity, the distance metric is based on Euclidean distance [1-4]. For two points, the distance simplifies to the absolute difference:

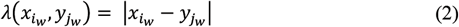

The distance cost *D* in DTW, which measures the similarity between two time series, is computed as the sum of distance values along the warping path of length *L*, which corresponds to the last cell of the cumulative distance cost matrix:

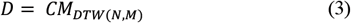

In our study, we employed a window size inspired by [2] to account for relevant time lags in the fMRI signals. We defined the window size based on the spectral characteristics of the activity signal, selecting a window with a −3*dB* cutoff at the low-frequency limit of *0*.*01HZ* for the fMRI signal [4, 8]. The window size *N* is calculated using:

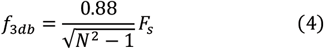

where *F*_*s*_ is the sampling frequency, and *f*_3*db*_ is the low cutoff frequency.

### B. DTW Normalization

For two signal pairs, the length of the warping path *L*, can vary. Since DTW distance is calculated as the sum of point-wise Euclidean distances along this path—represented by the final value in the distance cost matrix—summing distances across differing *L* values for signal pairs of the same length can lead to inconsistent DTW distance comparisons. To resolve this, we propose normalizing the DTW distance by the warping path length, improving its applicability in functional connectivity studies.

### C. DTW as an FNC metric

Since DTW is not upper-bounded, its distances are meaningful only in relation to other DTW values, emphasizing relative differences. To improve interpretability and enable comparison with traditional similarity measures like correlation, a transformation was proposed: first, the nDTW distance distribution across node pairs and subjects is demeaned, and then the values are multiplied by −1 [2]. Through this transformation, positive values represent strong similarity, negative values to indicate strong dissimilarity, and values near zero to reflect weak or irrelevant similarity. We apply this approach to both the DTW and nDTW measures.

### D. fMRI data & processing

We used resting-state fMRI data from 800 subjects provided by the Human Connectome Project’s (HCP) second session scans and the clinical dataset from the Function Biomedical Informatics Research Network (fBIRN) study. The HCP scans featured a repetition time (TR) of 0.72s, 72 slices, an echo time (TE) of 58ms, and a flip angle of 90°, with a voxel size of 2 × 2 × 2 mm, accumulating 1200 time points per subject. The fBIRN scans feature a TR of 2s, a total of 160 CN (age ± sd = 37.04 ± 10.86 years; 45 female / 115 male), and 151 SZ (age ± sd = 38.77 ± 11.63 years; 36 female / 115 male).

Following standard preprocessing—slice timing correction, realignment, spatial normalization, and smoothing [9] —we applied the NeuroMark ICA pipeline [10] to extract 53 intrinsic connectivity networks (ICNs). To capture relevant BOLD signal fluctuations, we used a Butterworth bandpass filter with a frequency cutoff of 0.01 − 0.15*HZ* [11]. The filter, with an optimal order of 7 as determined by MATLAB’s Buttord function, maintained a passband ripple under 3*dB* and a minimum stopband attenuation of 30*dB*. Finally, we standardized the filtered ICNs through z-score transformation.

For the HCP data, with *F*_s_ ≈ 1.39*Hz* (1/0.72 seconds), this results in a window size of roughly 123 time points. While the fBIRN data, with *F*_s_ ≈ 0.5*Hz* (1/2 seconds) results in a window size of 45 time points.

### E. Group analysis

We performed a Wald test on the data to compare CN with SZ using a generalized linear model to control for confounding factors including age, sex, site, and head motion [12, 13]. We corrected for multiple comparisons using the Benjamini-Hochberg false discovery rate (FDR) technique and we used Cohen’s *d* as an indicator of effect size. Furthermore, to characterize the sensitivity of the method in capturing group differences, the McNemar’s test is used. The nDTW is compared to classic DTW and Pearson correlation.

### F. Null hypothesis

The statistical validity of DTW as a functional connectivity measure is yet untested. Common approaches for generating fMRI surrogate data include phase randomization (PR) and autoregressive models, with PR often preferred for preserving data characteristics [14]. PR maintains signal correlation and captures coupling properties beyond linear correlations, though it risks rejecting the null hypothesis due to internal nonlinearities rather than inter-signal coupling [15].

To address this, time-shifted surrogate data (TS) was proposed, assuming no nonlinear interdependencies or significant cross-correlation between signals, though it requires a large enough shift to ensure distinct surrogates [15]. With the HCP dataset’s 1200 time points, this risk is reduced, allowing TS to complement PR in testing DTW’s sensitivity to nonlinear dependencies. If DTW rejects both TS and PR, the PR rejection likely reflects inter-signal dependencies beyond linear correlations rather than internal nonlinearities. Null models are constructed with functional connectivity networks (FNCs) from 1,000 surrogates per HCP subject and compared to original FNCs using the Wilcoxon rank-sum test [15] across both DTW and nDTW, under both TS and PR methods.

## III. Results

### A. Group analysis

Figure 1 illustrates the group comparison between CN and SZ in the fBIRN dataset. The upper triangles show that normalized DTW identifies more significant network pairs than both standard DTW and correlation.

**Figure 1.**
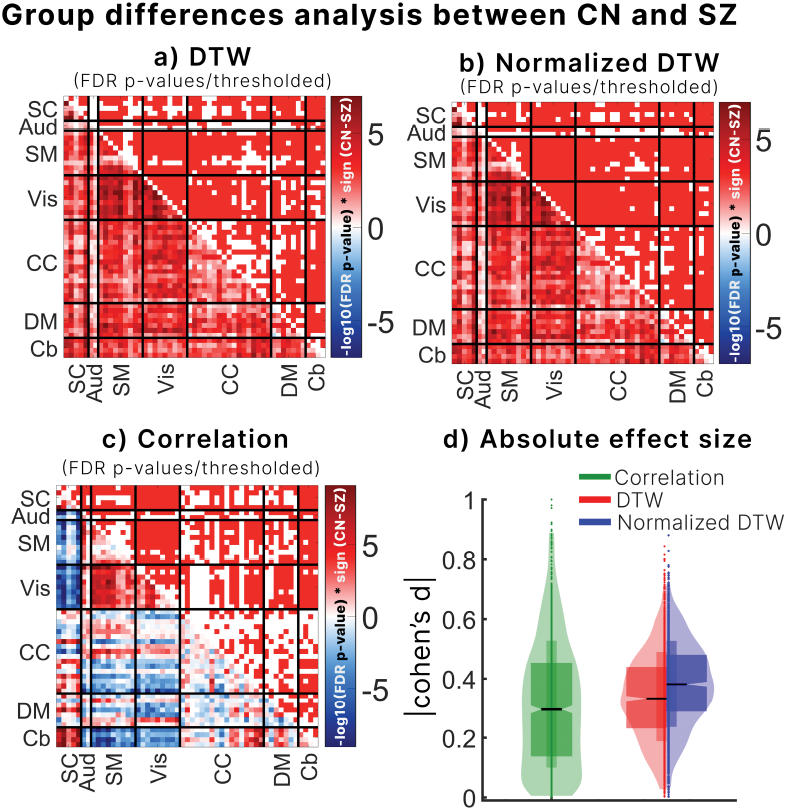
Compares schizophrenia patients with controls using standard DTW (a), normalized DTW (b), and correlation (c). The lower triangles display −log10 of FDR-corrected p-values, while the upper triangles show thresholded significant brain network pairs (p < 0.05). Normalized DTW detects the most significant differences, followed by standard DTW and correlation. Boxplots in the bottom right illustrate absolute effect sizes, with normalized DTW yielding the largest effect sizes, indicating it is the most sensitive to differences between groups.

Normalizing DTW distances by the warping path length enhances DTW’s clinical sensitivity (Figure 2), with significantly more exclusive network pairs identified by nDTW than by DTW (*p* < 0.0001; odds ratio = 55.33, McNemar’s test). Overlapping pairs identified by both DTW methods appear in the left panel of Figure 2, though normalized DTW (right-blue boxplot) yields higher effect sizes than standard DTW (left-red boxplot), underscoring its advantage.

**Figure 2.**
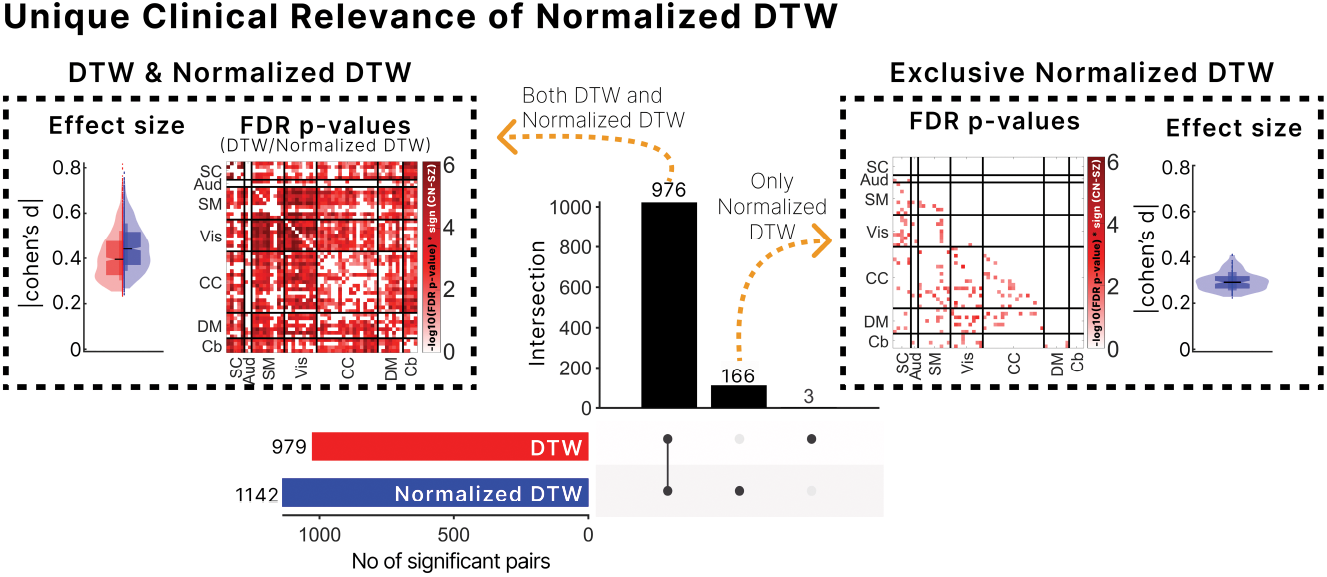
The clinical significance of normalized and standard DTW depicted using an UpSet plot. Nearly all brain network pairs detected by DTW are also identified by n DTW, with the latter capturing 166 additional pairs. This demonstrates that nDTW increases sensitivity without losing significance. On the right, pairs exclusive to normalized DTW are shown with effect sizes, while on the left, pairs identified by both DTW variants highlight the higher effect sizes from normalized DTW. Overall, normalized DTW detects more significant pairs compared to correlation, enhancing its clinical utility.

As shown in Figure 3b’s UpSet plot, nDTW identifies 1,142 clinically relevant pairs out of 1,378 unique brain network pairs, surpassing DTW’s 979 pairs and correlation’s 786 pairs. nDTW also demonstrates greater statistical sensitivity to CN vs. SZ differences than correlation (*p* < 0.0001; odds ratio = 4.33, McNemar’s test) and achieves higher effect sizes than both DTW and correlation (Figure 1d).

**Figure 3a.**
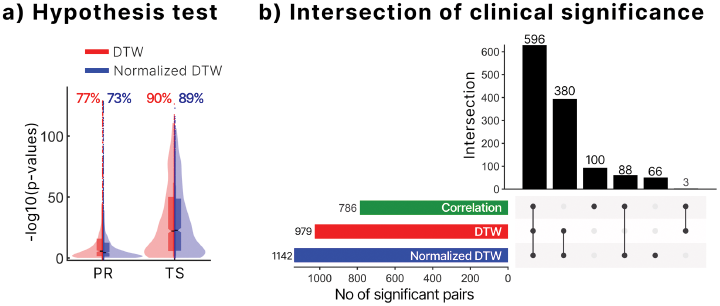
shows null hypothesis testing results using time-shifted and phase-randomized surrogates. Both standard and normalized DTW reject the null for many brain network pairs, as indicated by percentages above the box plots. Although standard DTW performs slightly better, rejection of both PR and TS surrogates suggests both metrics capture interdependencies beyond linear correlations. Figure 3b illustrates the overlap in significant network pairs detected by correlation, standard DTW, and normalized DTW. Normalized DTW identifies the most significant group differences, capturing 88 significant pairs that standard DTW misses, underscoring its enhanced sensitivity.

### B. Null hypothesis

The null hypothesis asserts that both DTW and nDTW reject the null for both TS and PR (Figure 3a), providing strong evidence that DTW captures interdependencies beyond linear correlations. Across both null models (TS and PR), DTW shows a slightly higher null rejection rate across features, suggesting it may be more sensitive to these interdependencies in the fMRI data than nDTW. However, this difference remains marginal.

## IV. Discussion

We propose a novel nDTW method as an FNC metric and compare its clinical sensitivity for SZ diagnosis with DTW and correlation approaches, while assessing its statistical relevance using PR surrogate data. In line with a previous study [1], our findings indicate that DTW-based metrics are more sensitive to clinical FNC alterations than correlation-based metrics. Notably, our proposed nDTW distances reveal greater differences between CN and SZ groups compared to DTW and correlation, highlighting nDTW as a superior tool for detecting psychosis-related brain changes.

Both DTW and nDTW reject null hypotheses generated from time-shifted and phase-randomized surrogates. The rejection of the PR null hypothesis, along with the rejection based on TS surrogates, provides strong evidence that DTW captures interdependencies beyond linear correlations. Designed to quantify similarities between time sequences with variable lags, stretches, and temporal deformations [5], the effective rejection of null models from TS and PR suggests that both DTW and nDTW distances successfully capture these complex interdependencies. Additionally, the DTW-derived warp elasticity method highlights the dynamics of stretching and shrinking in fMRI brain network time course relationships [4], underscoring the importance of considering interdependencies beyond linear correlations in functional connectivity studies [16, 17].

As illustrated in the upset plot in Figure 3b, nDTW identifies all but three significant network pairs detected by DTW. This indicates that the results captured by DTW are almost entirely encompassed within the normalized DTW, demonstrating that nDTW surpasses standard DTW in identifying significant group differences without missing any critical information. Furthermore, the effect size of the significant brain network overlaps between DTW and nDTW is higher for the latter, as shown by the blue box plot in Figure 2’s left panel. This implies that the nDTW not only retains almost all significant findings from DTW but also enhances the sensitivity to these differences.

Notably, nDTW captures clinical significance in 88 brain networks detected by correlation-based FNC but missed by the DTW. This improved detection may stem from nDTW’s slightly reduced sensitivity to interdependencies beyond linear correlation, allowing it to capture additional significances identified by linear measures.

While previous studies have emphasized DTW’s superiority over correlation in detecting clinical differences [1], test-retest reliability [1, 2], robustness to global signal regression [1, 2], and noise sensitivity [2], it is important not to misconstrue DTW or the nDTW as a replacement for correlation in functional network connectivity (FNC) analysis. As shown in Figure 3b, although correlation identifies fewer clinically significant network pairs than DTW or normalized DTW, it uniquely captures 100 significant brain network pairs. This underscores the complementary nature of correlation and DTW, though DTW—and especially nDTW—offer powerful advantages.

## V. Ethical standards & code availability

This study was conducted retrospectively using data collected from human subjects in compliance with all relevant ethical standards. The codes for all our analyses in MATLAB can be accessed through https://github.com/Sirlord-Sen/normalized_DTW_null_hypothesis.

## Acknowledgment

This work was supported in part by the NIH R01MH123610 and NSF 2112455.

## References

[1] A. C. Linke et al., “Dynamic time warping outperforms Pearson correlation in detecting atypical functional connectivity in autism spectrum disorders,” NeuroImage, vol. 223, p. 117383, 2020/12/01/ 2020, doi: 10.1016/j.neuroimage.2020.117383.

[2] R. J. Meszlényi, P. Hermann, K. Buza, V. Gál, and Z. Vidnyánszky, “Resting State fMRI Functional Connectivity Analysis Using Dynamic Time Warping,” (in English), Frontiers in Neuroscience, Methods vol. 11, 2017-February-17 2017, doi: 10.3389/fnins.2017.00075.

[3] S.-L. Wiafe, A. Faghiri, Z. Fu, R. Miller, and V. Calhoun, Capturing Stretching and Shrinking of Inter-Network Temporal Coupling in FMRI Via WARP Elasticity. 2024, pp. 1–4.

[4] S.-L. Wiafe, A. Faghiri, Z. Fu, R. Miller, A. Preda, and V. D. Calhoun, “The dynamics of dynamic time warping in fMRI data: a method to capture inter-network stretching and shrinking via warp elasticity,” Imaging Neuroscience, 2024, doi: 10.1162/imag_a_00187.

[5] K. K. Paliwal, A. Agarwal, and S. S. Sinha, “A modification over Sakoe and Chiba’s dynamic time warping algorithm for isolated word recognition,” Signal Processing, vol. 4, no. 4, pp. 329–333, 1982/07/01/ 1982, doi: 10.1016/0165-1684(82)90009-3.

[6] H. Sakoe and S. Chiba, “Dynamic programming algorithm optimization for spoken word recognition,” IEEE Transactions on Acoustics, Speech, and Signal Processing, vol. 26, no. 1, pp. 43–49, 1978, doi: 10.1109/TASSP.1978.1163055.

[7] C. A. Ratanamahatana and E. Keogh, “Everything you know about dynamic time warping is wrong,” in Third workshop on mining temporal and sequential data, 2004, vol. 32: Citeseer.

[8] S.-L. Wiafe, N. O. Asante, V. D. Calhoun, and A. Faghiri, “Studying time-resolved functional connectivity via communication theory: on the complementary nature of phase synchronization and sliding window Pearson correlation,” bioRxiv, p. 2024.06. 12.598720, 2024.

[9] W. D. Penny, K. J. Friston, J. T. Ashburner, S. J. Kiebel, and T. E. Nichols, Statistical parametric mapping: the analysis of functional brain images. Elsevier, 2011.

[10] Y. Du et al., “NeuroMark: An automated and adaptive ICA based pipeline to identify reproducible fMRI markers of brain disorders,” (in eng), Neuroimage Clin, vol. 28, p. 102375, 2020, doi: 10.1016/j.nicl.2020.102375.

[11] O. Josephs and R. N. Henson, “Event-related functional magnetic resonance imaging: modelling, inference and optimization,” Philosophical transactions of the royal society of london. series b: biological sciences, vol. 354, no. 1387, pp. 1215–1228, 1999.

[12] A. Yamashita et al., “Harmonization of resting-state functional MRI data across multiple imaging sites via the separation of site differences into sampling bias and measurement bias,” PLoS biology, vol. 17, no. 4, p. e3000042, 2019.

[13] A. Rao, J. M. Monteiro, J. Mourao-Miranda, and A. s. D. Initiative, “Predictive modelling using neuroimaging data in the presence of confounds,” NeuroImage, vol. 150, pp. 23–49, 2017.

[14] R. Liégeois, B. T. T. Yeo, and D. Van De Ville, “Interpreting null models of resting-state functional MRI dynamics: not throwing the model out with the hypothesis,” NeuroImage, vol. 243, p. 118518, 2021/11/01/ 2021, doi: 10.1016/j.neuroimage.2021.118518.

[15] G. Lancaster, D. Iatsenko, A. Pidde, V. Ticcinelli, and A. Stefanovska, “Surrogate data for hypothesis testing of physical systems,” Physics Reports, vol. 748, pp. 1–60, 2018/07/18/ 2018, doi: 10.1016/j.physrep.2018.06.001.

[16] A. Iraji et al., “The Nonlinear Brain: Towards Uncovering Hidden Brain Networks Using Explicitly Nonlinear Functional Interaction,” in 2023 IEEE 20th International Symposium on Biomedical Imaging (ISBI), 18-21 April 2023 2023, pp. 1–4, doi: 10.1109/ISBI53787.2023.10230347.

[17] S. Kinsey et al., “Networks extracted from nonlinear fMRI connectivity exhibit unique spatial variation and enhanced sensitivity to differences between individuals with schizophrenia and controls,” Nature Mental Health, vol. 2, no. 12, pp. 1464–1475, 2024/12/01 2024, doi: 10.1038/s44220-024-00341-y.

